# Serum C-terminal agrin fragment as a biomarker of age-related neuromuscular decline in men: influence of physical activity

**DOI:** 10.64898/2026.09.02.747348

**Authors:** Rami Hammad, Vincent Marcangeli, Marina Cefis, Rémi Chaney, Krystel Desjardins, Jordan Granet, Pierrette Gaudreau, Richard Robitaille, José A. Morais, Mylène Aubertin-Leheudre, Marc Bélanger, Gilles Gouspillou

**Author notes:** **Corresponding author :** Gilles Gouspillou : Département des sciences de l’activité physique, Faculté des sciences, Université du Québec à Montréal (UQÀM), Pavillon des sciences biologiques (SB), Local: SB-4640. 141, Avenue du Président Kennedy, Montréal, Québec, Canada, H2X 1Y4.

## Abstract

**Background:** Aging is associated with progressive neuromuscular junction (NMJ) degeneration. The C-terminal agrin fragment (CAF), a circulating product of agrin cleavage, has been proposed as a minimally invasive biomarker of NMJ degradation, but its relationships with specific neuromuscular and functional characteristics, and the potential modulating role of physical activity, remain poorly defined.

**Objective:** To examine age- and physical activity-related differences in serum CAF and to test its associations with neuromuscular integrity and functional capacity across the adult male lifespan.

**Methods:** We studied 136 community-dwelling men aged 20-92 years (active, n=87; inactive, n=49). Serum CAF was measured by ELISA; NCAM-positive fiber proportion, a widely used marker of myofiber denervation, was assessed by immunolabeling of vastus lateralis biopsies (n=127). Neuromuscular assessments included maximal voluntary isometric contraction (MVIC), lower-limb muscle power, electromechanical delay (EMD), nerve conduction velocities, H/M ratio, central activation ratio, co-activation, and twitch characteristics. Functional capacity was assessed with the 6-minute walk test, alternating step test, Timed Up and Go, sit-to-stand tests, and fast gait speed.

**Results:** Serum CAF increased progressively with age (β= 21,51, p<0.001) and was positively associated with NCAM-positive fiber proportion (β= 119,5 p=0.01). Higher CAF was associated with lower MVIC (β=-0.01, p=0.026), reduced lower-limb power (β= -0.01, p=0.027), and poorer performance on the 6MWT, alternating step test, TUG, sit-to-stand, and gait speed (p<0.05). CAF was associated with longer EMD in the whole cohort (β= 0.003, p=0.017) and inactive men (β= 0.005, p=0.007), but not in active men, and with lower H/M ratio in inactive men only (β= -0.003, p=0.0217). No associations were found with nerve conduction velocities, central activation ratio, co-activation, or twitch amplitude. Active men showed lower CAF than inactive men, particularly after age 70 (p=0.023).

**Conclusion:** In community-dwelling men, serum CAF is associated with age-related declines in neuromuscular and functional capacity and shows tissue-level consistency with the proportion of NCAM-positive fibers, supporting its use as an accessible biomarker of NMJ remodeling. Its limited association with most electrophysiological measures indicates it captures a specific component of neuromuscular aging rather than a global index of neuromuscular function. Physical activity was associated with a more favorable CAF profile, particularly among older men, identifying physical activity as a potential modifiable factor in age-related NMJ decline.

**Highlights:**

- Serum CAF rises progressively across the adult lifespan in men.
- CAF levels correlate with muscle fiber denervation markers.
- Elevated CAF is linked to poorer neuromuscular and physical function.
- Physical activity is associated with a more favorable CAF profile.

## Introduction

Aging is associated with a progressive decline in skeletal muscle strength and functional capacity, increasing the risk of mobility limitations, falls, and dependence [1–4]. Importantly, the age-related reduction in muscle strength often exceeds the loss of muscle mass, indicating that factors beyond muscle atrophy contribute substantially to impaired force production in older adults [5,6].

Among these factors, deteriorations of the neuromuscular system are thought to play a central role. Aging is associated with motoneuron loss, incomplete reinnervation of denervated muscle fibers, remodeling of motor units, and impaired neuromuscular junction (NMJ) integrity [7–12]. These changes may contribute to reduced voluntary activation, altered spinal excitability, slower nerve conduction, increased antagonist muscle coactivation, and ultimately lead to a poorer neuromuscular performance [13–16]. However, direct assessment of neuromuscular junction degeneration in humans remains challenging, creating a need for accessible biomarkers that reflect neuromuscular integrity and its functionality consequences.

Neural cell adhesion molecule (NCAM) positive fibers have commonly been used as an index of muscle fiber denervation, as NCAM expression is increased in denervated fibers and has been reported in aging and muscle disuse [17–24]. However, NCAM expression is not consistently elevated in all contexts of denervation, raising concerns about its specificity and reliability as a standalone marker of age-related neuromuscular degeneration [9]. Moreover, because NCAM assessment requires muscle tissue, it is not easily scalable for larger human studies or clinical screening. These limitations highlight the need for complementary circulating biomarkers of NMJ integrity.

Agrin is a particularly interesting candidate in this context because it plays a central role in the formation and maintenance of the NMJ. Neural agrin, released by motor neurons in the synaptic basal lamina, binds to the low-density lipoprotein receptor-related protein 4 and activates muscle-specific kinases. This pathway drives the clustering and stabilization of acetylcholine receptors at the postsynaptic membrane, thereby indicating efficient neuromuscular transmission and synaptic integrity [25]. Disruption of agrin signaling, or excessive degradation of synaptic agrin, may therefore compromise acetylcholine receptor organization and effective NMJ communication.

The C-terminal agrin fragment (CAF), is generated by the proteolytic cleavage of agrin, mainlyby synaptic neurotrypsin, and because this fragment can be detected in blood, circulating CAF has been proposed as a minimally invasive biomarker of NMJ remodeling or degradation [26]. Elevated CAF levels have been reported in aging, sarcopenia, cachexia, and neuromuscular disorders, and have been associated with muscle dysfunction in several clinical contexts [26–31]. Thus, CAF represents an attractive biomarker as it is directly linked to a molecular pathway required for NMJ functional integrity, while being measurable through a simple blood test.

However, despite this strong biological rationale, studies that have assessed the usefulness of CAF as a biomarker of neuromuscular function across from adulthood to late age remain scarce [32]. In addition, whether circulating CAF reliably tracks age-related neuromuscular impairment and functional decline, and whether habitual physical activity modifies this trajectory remain largely unresolved. Establishing whether CAF is associated with neuromuscular performance and neurophysiological determinants of force production is therefore essential before it can be considered a meaningful biomarker of NMJ integrity across the lifespan.

Thus, the present study aimed to assess whether immunoreactive circulating CAF levels is a relevant biomarker of age-related neuromuscular functional decline. Specifically, we examined age- and habitual physical activity-related changes in CAF levels, and investigated its associations with functional capacity, neuromuscular performance, and selected neurophysiological variables involved in muscle force production such as central activation ratio, spinal cord excitability, nerve conduction velocity and electromechanically delay.

## Material and Methods

### Participants

This cross-sectional study was tied to our previous publication [33] in which a total of 139 men were recruited. In the present manuscript, data from 136 community-dwelling men, aged 20 to 92 years, were included and analyzed. Participants were divided into two groups based on their physical activity status (active n=87 or inactive n=49) as well as four age groups (n=43: 20-39, n= 38: 40-59, n= 26: 60-69 and n= 29: 70-92 years old). All procedures were approved by the Comité Institutionnel d’Éthique de la Recherche avec des Êtres Humains (CIÉR #2020-2703) at Université du Québec à Montréal (UQAM). Informed consent was obtained from each participant after a whole explanation of the potential risks prior the start of data collection. All participants had to meet the following inclusion/exclusion criteria: body mass index between 18 and 35 kg/m², non-smokers (self-reported), limited alcohol intake (<2 drinks/day), and no history of motor, cardiac, neurological or psychological disorders.

### Physical Activity Status

To be considered active, participants had to meet at least one of the following criteria (see [33] for more details): i) engage in a minimum of 150 min per week of structured physical activity of moderate to high intensity; ii) achieve a daily step count of at least 10,000 steps; or iii) maintain a metabolic equivalent of daily tasks (MET) ≥ 1.6.

Briefly, criteria 1 was obtained using a self-reported physical activity obtained through a structured interview with a trained kinesiologist. Questions asked during the interview were derived from the validated questionnaire physical Activity Scale for the Elderly (PASE) [34]. Participants were asked to report their usual physical activity representative of the past 5 years. Criteria 2 and 3 (objective measures) were assessed using tri-axial accelerometer (SenseWearMini Armband) [35]: Participants were asked to wear the armband on the non-dominant arm for at least 3 days and ideally for 7 days. Data were taken from days when the armband was worn at least 80% of the time over 24h (>19h and 12 min). Data were analyzed using the software Armband Sensewear 8.1.

### Anthropometric Data and Body Composition

Height was measured (cm) using a wall-mounted height gauge (SECA 67029, Hanover, MD, USA) and mass (kg), with an electronic scale (ADAM - GFK 660a, USA). The BMI was calculated using the following formula: Mass (kg)/Height² (m²) [36].

DXA (GE Prodigy Lunar) was used to assess total, relative and lower limb lean masses. Participants were fasting for a minimum of 2 h before the DXA scan.

### Assessment of functional capacity

1. 6-minute walk test (6MWT): participants were instructed to walk as far as possible within 6 minutes without receiving verbal encouragement. The test was conducted on a 25-m track. Total distance covered was recorded in meters [37].
2. Alternate step test: Participants stood facing a 20-cm-high step and were instructed to alternately place their right and left foot on the step as quickly as possible for 20 seconds. The total number of steps completed was recorded [38,39].
3. Sit-to-stand test: Participants were asked to stand up and sit down repeatedly from a chair as quickly as possible for 30 seconds, with arms crossed over the chest. The time required to complete 5 and 10 repetitions and the total number of repetitions performed within 30 seconds were recorded and analyzed [40].
4. Timed Up and Go (TUG) test: Participants were instructed to stand up from a chair, walk a distance of 3 meters, turn, walk back, and sit down as quickly as possible. The total time to complete the task was recorded [41,42].
5. 4-meter fast walk test: Participants were instructed to walk at their fastest safe walking speed along a straight walkway. A pre-measurement acceleration distance was provided before the 4-meter timed section to allow participants to reach their maximal walking speed before timing began. Timing started when the participant’s first foot crossed the beginning of the 4-meter marked section and stopped when the first foot crossed the 4-meter endpoint. The time required to complete the 4-meter distance was recorded, and walking speed was calculated as meters per second (m·s ¹) [43].

### Assessment of Muscle Strength and Power

Maximal voluntary isometric contraction strength of the knee extensors (MVIC-E) and flexors (MVIC-F) were assessed on the right lower limb on a modified BTE chair (Primus RSChair, BTE). A small platform, set at 135° with respect to the seat, was fixed to the front of the seat at one end to an analog strain gauge at the other end. The participants were seated with the knee and hip joint angles fixed at 135^○^ and 90^○^, respectively, and the tested (right) leg secured to the platform at the level of the malleoli. Participants were instructed to contract as quickly and forcefully as possible and to maintain the contraction until prompted to relax (≈3 s MVIC). Maximal force (extension; MVIC_E), rate of force development (RFD; up-slope), and rate of force relaxation (RFR; down-slope) were determined from the force curves of muscle extension contraction.

Lower limb muscle power (LLMP) was measured on the right lower limb from a seated position using the Nottingham Leg Extensor Power rig (University of Nottingham, UK). Participants were instructed to push the pedal as hard and fast as possible, accelerating a flywheel attached to an analog to digital converter and computer. Several (3-5) trials were recorded for each participant for both force and power measurements, and the best one was selected for analysis.

### Assessment of Muscle Twitch Characteristics and Central Activation Ratio

From the same system and setup as for the MVIC test, two self-adhesive electrodes (5×9 cm) were placed on each side of the central area (muscle belly) of the right Vastus Lateralis (VL). With the knee extensor muscle relaxed, three supramaximal electrical bursts were randomly elicited using a 5 ms stimulation train (1 ms square pulse delivered at 500 Hz) (Grass S88 stimulator and a SIU5 isolation unit; Astro-med Inc., West Warwick, RI, USA) at an interval ranging between 15 and 20s. The stimulation intensity was progressively increased until a maximal evoked response was obtained and then further increased to ensure supramaximal stimulation. These supramaximal evoked responses were recorded to determine muscle twitch amplitude, twitch contraction time (CT), half-relaxation time (½RT), and twitch electromechanical delay (EMD). To assess the central activation ratio (CAR), 5 ms stimulation train (1 ms square pulse delivered at 500 Hz) were elicited and superimposed (SIT) mid contraction (≈ 1.5 s) during 3 additional 3 s MVIC-E separated by 1 min rest to assess an individual’s ability to fully activate the muscles, providing an estimate of CAR [CAR (%) = MVIC_E force / (MVIC_E + SIT force) * 100 or strength reserve (SR) [100-CAR].

### Electromyographic (EMG) Recording

The skin surfaces overlying the right Soleus (Sol), VL and Biceps Femoris short head (BF) muscles and the tibial nerve in the popliteal fossa, were prepared by shaving, gently rubbing (18.1 mm 3M™ Red Dot™ Trace Prep, 2236, 3M, St. Paul, MN 55144-1000, U.S.A.) to remove dead or dry skin, and clean with pre-soaked 70% v/v isopropyl alcohol swabs (Canadian Custom Packaging Company, Toronto, Canada) for disinfection. Two electrodes (hypoallergenic conductive adhesive hydrogel foam electrodes; 8 mm; Model Medi-Trace 133, Covidien Kendall MA, USA), with an inter-electrode distance approximately 10 mm, were placed over the Sol below the medial Gastrocnemius muscle. For the VL EMG, two other electrodes were placed near the muscle belly. Two other electrodes were placed just superior of the lateral border of the popliteal fossa to record the BF. A ground electrode was placed overlying the mid-anterior surface of the tibia. The EMG signals were amplified (Grass P511; Astro-Med Inc., West Warwick, RI, USA) and bandpass filtered (10-300 Hz) before being digitized at 10 kHz on a 12-bit data acquisition system (Axoscope 10 and Digidata 1440A; Molecular Devices, Sunnyvale, CA, USA). The recording apparatus were connected to a UPS-isolation system to ensure a continuous power without unwanted spikes. EMG was used to obtain the electromechanical delay (EMD) during the MVIC-E (the latencies between the onset of the activation and the start of force development), the cross-correlation muscle activation between the agonist (knee extensors) and antagonist (knee flexors) muscles during the MVIC.

### Soleus Hoffmann (H) reflex and Motor (M) Wave Measurements

With the participants lying prone on a cushioned table, with the arms and shoulders relaxed and the head aligned along the axis of the neck, progressively increasing 1 ms pulses were randomly applied every 5-8 s to the tibial nerve in the popliteal fossa. The Sol EMG amplitude was obtained using the Clampfit software (Molecular Devices, USA) and allow for the production of a H-M recruitment curve and the calculation of the Hmax/Mmax [maximal H (Hmax)/maximal M (Mmax) ratio. The Hmax/Mmax is used as an indicator of spinal cord excitability.

The latencies of the M and H responses were obtained from the Sol EMG signal. An estimate of both the motor neurons and the Ia afferent neurons conduction velocities (MN_CV_ & Ia_CV_) was calculated, for more details see [44].

### Quantification of circulating serum C-Terminal Agrin Fragment

Blood serum was isolated from 5 mL venous blood samples obtained from the median cubital vein of all participants. Serum C-terminal agrin fragment (CAF) immunoreactive levels were measured using anenzyme-linked immunosorbent assay (ELISA) kit (Human Agrin SimpleStep ELISA, Ab216945; Abcam, Cambridge, UK). Samples were diluted 1:4 and analyzed in duplicate following the manufacturer’s protocol.

### In situ immunolabelling for Neural Cell Adhesion Molecule

Vastus lateralis muscle samples were mounted in tragacanth, frozen in cooled liquid isopentane, and stored at -80°C. Serial cross sections of 10 µm thickness were cut with a cryostat at -20°C and mounted on lysine coated slides (Superfrost) to assess the proportion of fibers positive for neural cell adhesion molecule (NCAM). Muscle cross sections were allowed to reach room temperature and rehydrated with phosphate-buffered saline (PBS) before being incubated for 30 minutes in a blocking solution with goat serum (10% in PBS 1X). Sections were then incubated for 2 hours at room temperature with mouse monoclonal IgG1 anti-NCAM (BD347740, 1/50) and mouse monoclonal IgG2b anti-dystrophin (MilliporeSigma, D8168, 1/500) primary antibodies. Sections were then washed 3 times in PBS 1X and then incubated for 1 hour with Alexa Fluor 594 goat anti-mouse IgG1 (Thermo Fisher Scientific, A-21125, 1/500) and Alexa Fluor 488 goat anti-mouse IgG2b (Thermo Fisher Scientific, A-21141, 1/500) secondary antibodies. Slides were then washed 3 times in PBS 1X, fixed with 5%formaldehyde for 12 minutes and finally coverslipped using Prolong Diamond (Thermo Fisher Scientific, P36961) as mounting medium. Slides were imaged using an Olympus IX83 Ultra Sonic fluorescence microscope (Olympus, Tokyo, Japan). The proportion of NCAM-positive fibers were manually quantified using ImageJ (NIH, Bethesda, MD, USA; https://imagej.nih.gov/ij/) and a minimum of 200 myofibers was analyzed per sample (810.4 in average; SEM = 34.71; SD = 386.5).

## Statistical Analyses

All statistical analyses were done using Prism statistical software (version 11.0.0). A significance level of p<0.05 was established a priori for all inferential tests. Participant’s demographic and anthropometric variables were characterized using descriptive statistics, presented as mean ± standard deviation (SD), accompanied by the range (minimum to maximum). The relationship between muscle function variables, neurophysiological variables and serum C-Terminal Agrin fragment levels was investigated using simple linear regression and one-tailed Pearson correlation coefficient methods.

These analyses aimed to quantify the linear associations between serum CAF concentrations and neuromuscular variables, as well as to characterize age-related patterns observed across the cross-sectional sample. To ascertain the influence of physical activity status on age-related physiological alterations, the slopes and intercepts from simple linear regression models were compared for active and inactive participants. Furthermore, the interaction between physical activity and age groups was examined using Two-way ANOVA to determine if the age-related changes of the neuromuscular function and serum C-Terminal Agrin fragment and NCAM variables differed significantly based on physical activity status and/or age groups.

## Results

### Participants’ characteristics

Table 1 presents the anthropometric characteristics of the 136 participants, including subgroup data for active and inactive men. These data have been reported previously in [33]. No significant differences were observed between active and inactive groups for age, body mass and height. Total lean mass was significantly higher in active participants while BMI was significantly lower in active participants (Table 1).

**Table 1:** Participants anthropometric characteristics.

| Variables | All (n=136) |  | Inactive (n=49) |  | Active (n=87) |  | P value |
| --- | --- | --- | --- | --- | --- | --- | --- |
| | Mean ± SD | Range | Mean ± SD | Range | Mean ± SD | Range | |
| Age (y) | 51.8 ± 18.6 | 20.0 – 92.0 | 54.4 ± 19.8 | 24.0 – 92.0 | 50.3 ± 17.8 | 20.0 – 84.0 | P = 0.219 |
| Body Mass (kg) | 79.4 ± 12.0 | 51.8 – 110.0 | 82.7 ± 12.6 | 54.8 – 110.0 | 77.4 ± 11.3 | 51.8 – 109.0 | P = 0.054 |
| Lean Mass (kg) | **56.4 ± 6.8** | **40.9 - 74.7** | **54.9 ± 6.7** | **42.7 - 72.0** | **57.2 ± 6.7** | **40.9 - 74.7** | **P = 0.012** |
| Height (m) | 1.74 ± 0.07 | 1.60 - 1.92 | 1.75 ± 0.08 | 1.60 - 1.90 | 1.74 ± 0.06 | 1.60 - 1.92 | P = 0.656 |
| BMI (kg/m<sup>2</sup>) | **26.1 ± 3.6** | **19.0 - 34.5** | **27.1 ± 3.7** | **19.6 - 33.3** | **25.5 ± 3.5** | **19.0 - 34.5** | **P = 0.013** |

The mean ± standard deviation (SD) and the range (minimum - maximum) values for key anthropometric variables for all participants and for the categorized inactive and active groups.

### The impact of aging and physical activity status on the percentage of NCAM-positive fibers and serum C-terminal Agrin Fragment (CAF) levels

This study investigated first the impact of aging and physical activity on the proportion of NCAM-positive fibers, a widely used marker of denervation, on skeletal muscle cross sections, using immunolabeling (Fig. 1A). Although minimal to no effect of aging or physical activity status could be evidenced when data were stratified by age groups (Fig. 1B), a progressive age-related increase in the proportion of NCAM-positive fibers was observed in the whole cohort (all participants combined) and in the inactive group, but not in active participants (Fig. 1C) when age was used as a continuous variable. Collectively, these data suggest an age-related increase in fibers with altered or denervated NMJ and suggest a potential protective impact of physical activity status.

**Figure 1:**
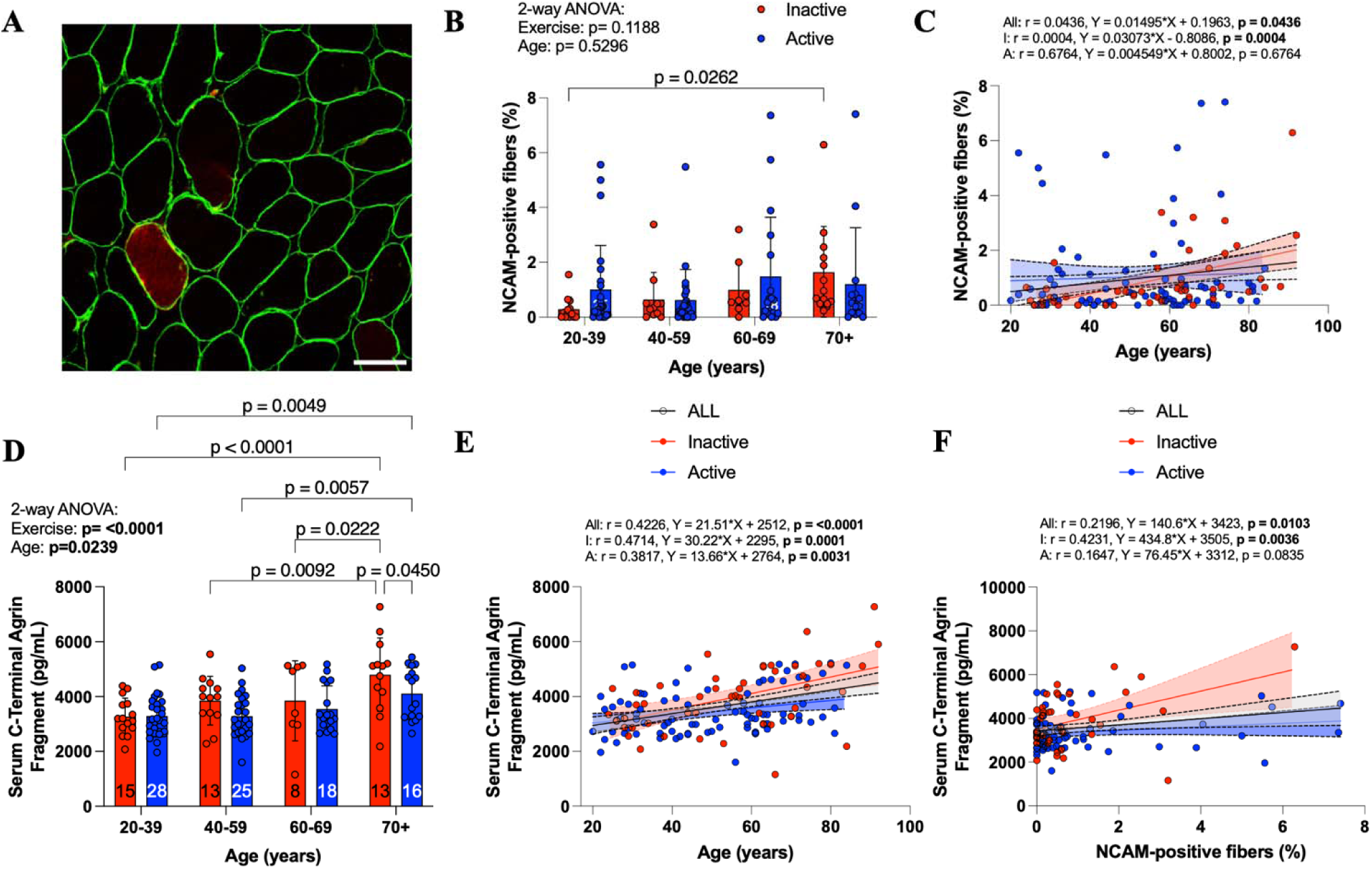
Impact of aging and physical activity status on the proportion of NCAM-positive fibers and the circulating level of CAF in men. (A) Representative immunolabeling for dystrophin (green) and NCAM (red) used to assess the proportion of NCAM positive fibers. (B) Quantification of the proportion of NCAM-positive fibers in inactive and active participants across age groups (n= 127: 20-39, 40-59, 60-69, and 70-92). (C) Relationship between age (expressed as a continuous variable) and the proportion of NCAM-positive fibers for all participants (n = 127) and stratified by activity status. (D) Serum CAF levels in inactive and active participants across age groups (n= 136: 20-39, 40-59, 60-69, and 70+). (E) Relationship between age (expressed as a continuous variable) and Serum CAF concentrations for all participants (n= 136) and stratified by activity status. (F) Relationship between the proportion of NCAM-positive fibers and Serum CAF levels in the whole cohort. Statistical significance was set at *p < 0.05*.

Then we investigated whether circulating levels of CAF mirrored changes in the proportion of NCAM-positive fibers in this cohort. As seen in Fig. 1D and 1E, the circulating levels of CAF progressively increase with age. Interestingly, when data were stratified by age groups, a significant protective effect of physical activity status was observed (Fig 1D), particularly evident in individuals aged 70 and over, on the circulating levels of CAF. Positive associations between the proportion of NCAM-positive fibers and circulating levels of CAF were observed in the present cohort (Fig 1F).

Taken altogether, our data indicate i) that aging in men is associated with alterations in NMJ integrity, ii) that physical activity may partly protect against age-related alterations in NMJ integrity and iii) strengthen available evidence positioning circulating CAF as a potentially useful biomarker of NMJ integrity status in humans.

### Serum CAF levels are associated with functional capacity in men

With the aim of assessing the clinical value of serum CAF to identify individuals with low physical performance and muscle health, this paper investigated whether this biomarker displayed associations with validated measures of muscle force, power and functional capacity. As shown in Figure 2, serum CAF levels were negatively associated with MVIC and lower limb muscle power in the whole cohort (Fig. 2A and B). Serum CAF levels were also negatively associated with participants’ performance at the 6-minute walk test, alternate step test, Timed Up and Go test and variations of the sit-to-stand test (Fig. 2C-H) in the whole cohort. Serum CAF concentrations were also negatively associated with participants’ walking speed in the full cohort (Fig. 2I). Interestingly, no striking difference in the aforementioned relationships could be observed between active and inactive participants (Fig 2). Taken altogether, these data demonstrate that circulating CAF is associated with muscle and physical performance in men, with higher levels being associated with lower muscle and physical performance.

**Figure 2.**
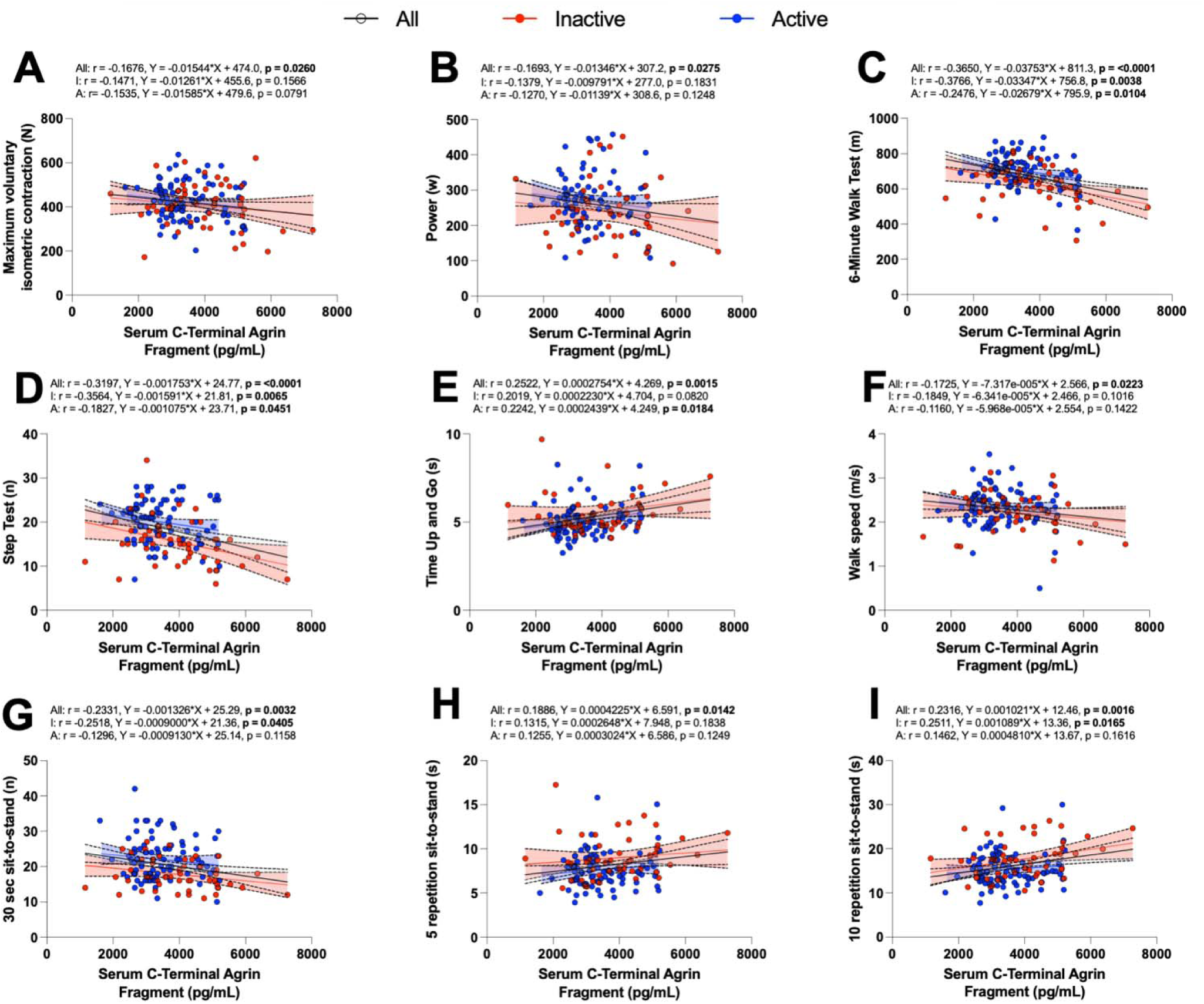
Associations between serum C-terminal Agrin Fragment (CAF) concentration levels and Force, Power and physical performance in physically active and inactive men. Scatter plots with linear regression lines and 95% confidence intervals depict the relationships between serum CAF levels and: (A) maximum voluntary isometric contraction (MVIC, N) and (B) lower limb muscle power (W), strat fied by physical activity status. **(C)** 6-minute walk test distance (6MWT), **(D)** alternating step test performance, **(E)** Timed Up and Go (TUG) duration, **(F)** fast gait speed, **(G)** 30-second sit-to-stand repetitions, **(H)** 5-repetition sit-to-stand time, and **(I)** 10-repetition sit-to-stand time. Data are shown for inactive (red circles-Inactive) and active (blue circles-Active) men, as well as for the overall cohort (black line). Regression coefficients (β) and p-values are reported for each group. Statistical significance was set at *p* < 0.05.

Serum CAF levels were nether significantly associated with central activation ratio (Fig. 3A), nor spinal cord excitability (H/M ratio) in the whole cohort or in active men; however, a significant negative association was observed in inactive men (Fig. 3B). Rates of force development, motor and sensory nerve conduction velocities, and co-activation were not significantly associated with serum CAF levels in the overall cohort or in the active and inactive subgroups (Fig. 3C-F). Neither absolute (N), nor relative (% MVIC force) twitch amplitude was significantly associated with serum CAF levels in the overall cohort or in each subgroup (Fig. 3G-H). In contrast, electromechanical delay increased significantly with higher serum CAF Levels in the whole cohort and in inactive men, whereas no significant association was observed in active men (Fig. 3I). In all, these findings indicate that electromechanical delay and, in inactive men, the H/M ratio were the only neuromuscular parameters significantly associated with serum CAF levels.

**Figure 3.**
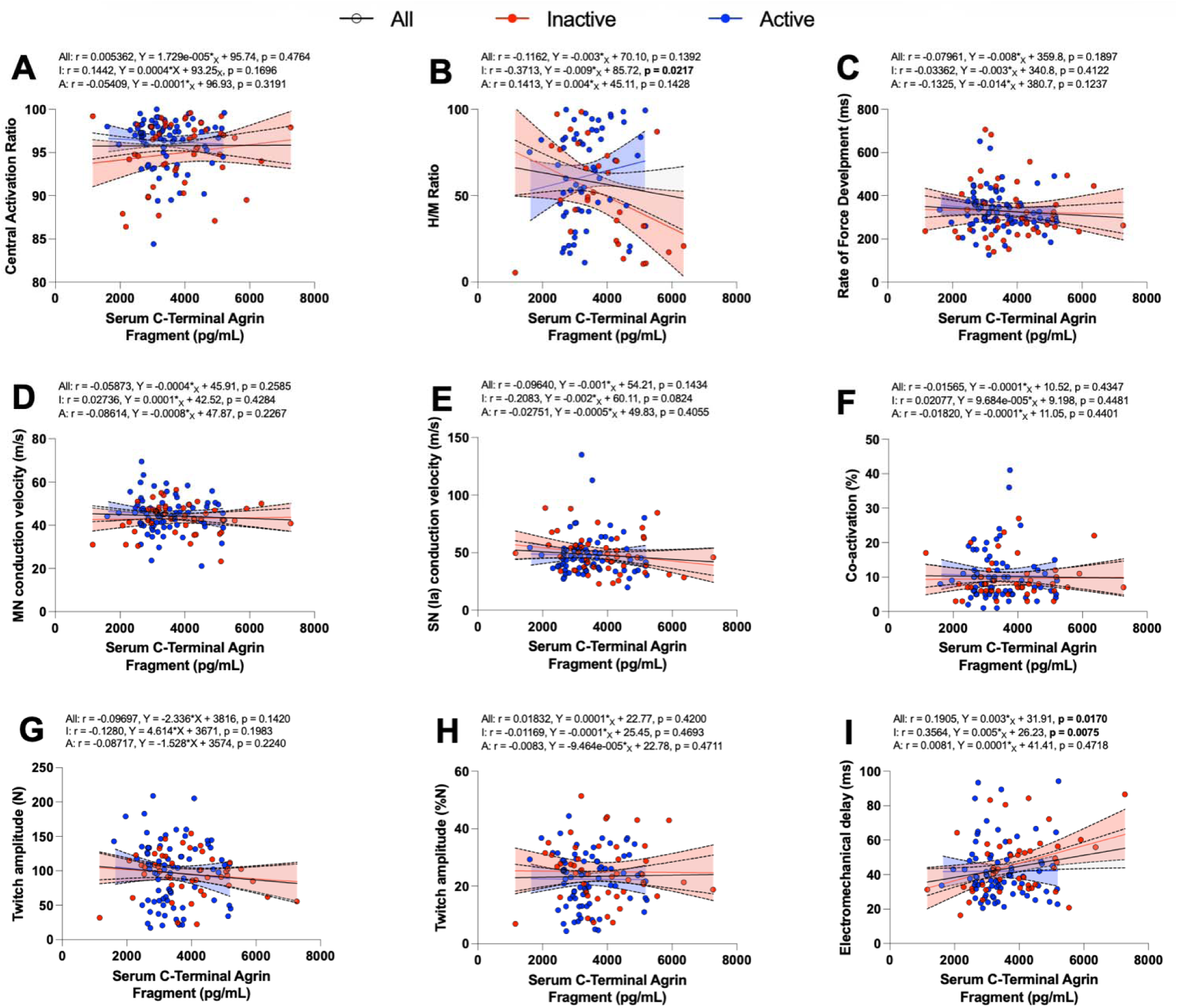
Association between serum C-terminal Agrin Fragment (CAF) levels and neuromuscular variables in active and inactive men. Scatter plots with linear regression lines and 95% confidence intervals depict the relationships between serum CAF (pg/mL) and (A) central activation ratio (%), (B) spinal cord excitability - H/M ratio, (C) rate of force development (ms), (D) motor nerve (MN) conduction velocity (m/s), (E) sensory nerve (SN) conduction velocity (m/s), (F) co-activation (%), (G) twitch amplitude (N), (H) twitch amplitude normalized to maximum force (%N), and (I) electromechanical delay (ms), for all participants (black line), inactive men (red circles), and active men (blue circles). Regression coefficients (β) and p-values are reported for each group. Statistical significance was set at *p < 0.05*.

## Discussion

Our understanding of the mechanisms driving age-related declines in functional capacity and mobility remains incomplete, and the potential beneficial role of habitual physical activity is often overlooked. Although degeneration of the motor unit and neuromuscular junction (NMJ) is increasingly recognized as a contributor to age-related neuromuscular decline, direct assessment of NMJ integrity in humans remains challenging. In this context, the present study investigated whether circulating C-terminal agrin fragment (CAF), a blood-based marker generated by agrin cleavage, is associated with age, physical activity status, muscle performance, functional capacity, and selected neurophysiological variables in men across the adult lifespan. The main findings were that serum CAF Levels increased with age, were generally lower in active than inactive individuals, and were consistently associated with lower force, power, and functional capacity. In addition, serum CAF levels were positively associated with the proportion of NCAM-immunoreactive fibers, supporting the validity of CAF as a circulating biomarker related to NMJ alteration. In contrast, associations between CAF and specific neuromuscular/electrophysiological variables were more limited, being restricted primarily to electromechanical delay in the whole cohort and inactive men, and to the spinal cord excitability (H/M ratio) in inactive men. Together, these findings support the relevance of CAF as an accessible biomarker associated with age-related neuromuscular and functional decline, while also indicating that CAF should be interpreted with cautions regarding various components of neuromuscular function.

In the present study, serum CAF increased progressively with age, consistent with previous reports of elevated circulating CAF in older adults and in populations with muscle wasting, weakness, or sarcopenia [27,29,32,45,46]. Because CAF is generated by proteolytic cleavage of agrin, a key organizer of acetylcholine receptor clustering and NMJ stability, higher circulating CAF is commonly interpreted as reflecting increased NMJ degeneration or remodeling [26,28]. Serum CAF was also positively associated with the proportion of NCAM-positive fibers, a marker upregulated in denervated or partially denervated fibers [17,21,23,24]. This tissue-level association strengthens the biological validity of CAF as a minimally invasive marker of neuromuscular alterations in humans.

Physical activity appeared to modulate age-related changes in CAF: active participants generally showed lower serum CAF than inactive individuals, an effect most apparent in older age groups. This aligns with evidence that habitual physical activity is associated with better preservation of motor unit structure, reinnervation capacity, and neuromuscular performance [12,21,23,47]. Conversely, sedentary behavior and prolonged bed rest have been associated with detrimental changes in older human motor unit properties [19,48,49]. The present findings further suggest that these neuromuscular benefits are accompanied by lower circulating CAF Levels in physically active men, consistent with findings in older dancers who exhibit lower CAF levels and superior balance and gait performance than sedentary peers [50]. However, activity-group differences were modest and not uniform across age categories or outcomes, suggesting that habitual physical activity may attenuate, but not prevent, age-related NMJ modifications. More specific modalities, such as resistance, power training or high intensity interval training, may be needed to more robustly influence NMJ-related outcomes than reported general physical activity status alone.

A strength of this study is the breadth of functional assessment. Serum CAF was consistently associated with poorer performance across several clinically relevant tests, including the 6-minute walk test, alternating step test, Timed Up and Go, sit-to-stand performance, and fast gait speed, and it was negatively associated with maximal voluntary isometric contraction and lower-limb muscle power in the whole cohort. These associations indicate that higher CAF identifies individuals with poorer muscle and physical performance, likely reflecting biological processes that contribute, alongside muscle mass, muscle quality, excitation-contraction coupling, tendon properties, and central neural factors, to reduced functional capacity.

Despite this consistent link to functional outcomes, CAF showed limited association with the specific neurophysiological variables assessed. Central activation ratio, rate of force development, motor and sensory nerve conduction velocities, co-activation, and absolute or normalized twitch amplitude were not significantly associated with serum CAF. The absence of associations between CAF and central activation ratio, H/M ratio of the inactive individuals, nerve conduction velocity, or co-activation suggests that circulating CAF is unlikely to reflect impairments in central neural activation, peripheral nerve conduction, or motor control. Instead, these findings indicate that CAF is more closely linked to peripheral mechanisms underlying muscle force generation. This interpretation is further supported by the observed association between CAF and electromechanical delay (EMD), which reflects processes involved in excitation–contraction coupling and force transmission. Instead, CAF likely captures a more specific component of neuromuscular aging, such as NMJ remodeling, denervation-reinnervation dynamics, or peripheral synaptic instability, that influences function without tracking every neurophysiological measure.

Among neuromuscular variables, electromechanical delay (EMD) showed the most robust association with CAF, increasing significantly with higher CAF in the whole cohort and in inactive men, but not in active men. EMD, the interval between muscle electrical activation and measurable force, is shaped by neuromuscular transmission, excitation-contraction coupling, muscle-tendon stiffness, and contractile properties [51,52]. Higher CAF may therefore mark individuals with slower translation of activation into force, particularly among inactive men; however, because EMD is not NMJ-specific, this should be read as an association with delayed force production rather than direct evidence of impaired neuromuscular transmission.

The negative association between CAF and the H/M ratio in inactive men is similarly noteworthy but warrants caution, since the H/M ratio indexes spinal reflex excitability rather than NMJ integrity directly. It may instead suggest that, in inactive individuals, higher CAF coexists with broader neuromuscular alterations, such as motor unit remodeling, altered afferent feedback, or changes in Ia afferent-motoneuron excitability [53]. Its absence in active men suggests physical activity may alter the relationship between peripheral NMJ-related remodeling and spinal reflex function but given the indirect nature of the H/M ratio and the subgroup-specific finding, this result is best considered hypothesis-generating.

The lack of association between CAF and the rate of force development or twitch amplitude is perhaps unsurprising given how many central and peripheral factors are involved in rapid force production and evoked contractile responses, including voluntary neural drive, motor unit recruitment and discharge rate, fiber-type distribution, tendon stiffness, and muscle architecture [54,55], as well as muscle mass and excitation-contraction coupling for evoked twitch. NMJ-related alteration, as reflected by circulating CAF, is therefore likely one contributor among others to these outcomes rather than their main determinant.

Our findings support circulating CAF as a promising, minimally invasive biomarker of neuromuscular aging and functional decline. Unlike biopsy-based approaches, CAF can be measured in blood, making it feasible for larger studies, longitudinal monitoring, and intervention trials. Its interpretation should nonetheless remain cautious: CAF is not a standalone diagnostic marker of NMJ integrity, nor a direct measure of denervation, spinal excitability, or transmission speed, but may be most useful within a broader biomarker and functional-assessment framework for identifying individuals at risk of accelerated neuromuscular aging or mobility decline. Longitudinal studies will be needed to determine whether elevated CAF predicts future functional deterioration, and whether different types of exercise or other interventions can reduce CAF or modify its relationship with functional performance.

## Limitations

Our study has some limitations. First, only male participants were included. This choice was made to reduce biological heterogeneity and because men have been proposed to exhibit a more linear trajectory of neuromuscular aging (Piasecki et al., 2024). However, whether the present findings can be generalized to women remains to be confirmed, particularly given the potential influence of sex hormones on neuromuscular aging. Second, the cross-sectional design limits causal interpretation. Although serum CAF was associated with age, physical activity status, functional capacity, and selected neuromuscular variables, longitudinal studies are required to determine whether elevated CAF predicts subsequent neuromuscular or functional decline. Third, the study sample consisted exclusively of relatively healthy, community-dwelling men. Even participants classified as inactive demonstrated relatively high physical performance, with many showing profiles consistent with successful aging [57]. This may have limited the ability to detect stronger associations between CAF and neuromuscular impairment, and the present findings may not extend to frail older adults or individuals with overt neuromuscular disease.

Fourth, physical activity status was based on a combination of accelerometry and self-reported habitual activity, but the study was not designed to isolate the effects of specific exercise modalities. Therefore, the relative contributions of endurance, resistance, and power-based training to CAF Levels and NMJ-related outcomes could not be determined.

Fifth, although the positive association between serum CAF and NCAM-positive fiber proportion supports the biological relevance of CAF as a circulating biomarker related to NMJ integrity, neither measure provides a direct structural or functional assessment of the NMJ. CAF should therefore be interpreted as an indirect biomarker of NMJ remodeling rather than a definitive measure of NMJ degeneration.

Sixth, although participants had no self-reported renal disease, we cannot fully exclude a contribution of normal age-related decline in glomerular filtration rate to serum CAF, as CAF has been shown to correlate with renal clearance markers in other populations [58].

Finally, the neurophysiological assessments performed in this study capture selected aspects of neuromuscular function but do not encompass all mechanisms involved in age-related force and functional decline, such as motor unit number, motor unit firing behavior, muscle architecture, tendon stiffness, or excitation–contraction coupling. Future longitudinal and interventional studies combining circulating biomarkers with more direct measures of motor unit and NMJ structure/function will be needed to clarify the mechanistic and clinical significance of CAF.

## Conclusions

Serum CAF increased with age and was associated with poorer force, power, and functional capacity in men across the adult lifespan. Its positive association with NCAM-positive fiber proportion provides tissue-level support for CAF as a circulating biomarker of NMJ integrity, while physical activity, particularly in older individuals, was associated with a more favorable CAF profile, suggesting that an active lifestyle may partially attenuate or delay age-related neuromuscular remodeling. The limited associations between CAF and specific neuromuscular variables indicate that CAF captures only part of the neuromuscular aging process. Overall, these findings support serum CAF as an accessible biomarker of NMJ remodeling, neuromuscular aging, and functional decline, while underscoring the need for longitudinal and interventional studies to clarify its mechanistic meaning and clinical utility.

## Acknowledgments

We are grateful to all participants who generously contributed their time and efforts to this study. We also thank Jill Vandermeerschen for her insightful input and discussions regarding statistical analyses. Language editing of the manuscript was assisted by generative AI tools (Copilot and Claude). The tool was used exclusively to improve clarity and readability. All scientific content was independently developed and verified by the authors.

## Data Availability

The data that support the findings of this study are available from the corresponding author upon reasonable request.

